# Unsaturation and Approximate Isotopic Homogeneity in Leaf Air Spaces

**DOI:** 10.1101/2024.09.30.610858

**Authors:** Spencer E. Collaviti, Hilary Stuart-Williams, Graham D. Farquhar, Lucas A. Cernusak, Diego A. Márquez

## Abstract

- We consider two assumptions of leaf isotope gas exchange measurements: that leaf air spaces are saturated with water vapour, and that this vapour is of a homogeneous isotopic composition. In particular, we consider whether these assumptions can concurrently hold and, if not, which assumption is preferable to retain.
- We present two methods using independent measurements of both leaf surfaces to consider these assumptions. The first method determines the isotopic inhomogeneity between the abaxial and adaxial evaporative sites when saturation is assumed. The second method determines the unsaturation in the abaxial and adaxial substomatal cavities when isotopic homogeneity is assumed. The methods are applied on *Gossypium hirsutum* (cotton) under benign atmospheric demand conditions (1.0 kPa air saturation deficit).
- We find evidence that assuming saturation contradicts isotopic homogeneity and vice-versa. We compare each assumption to pre-existing data and find that it is reasonable to assume isotopic homogeneity, but not leaf vapour saturation. Thus, we find that leaves experience unsaturation even under benign atmospheric demand conditions and have a spatial variation in their unsaturation, with lower humidities associated with the surface of least stomatal resistance.
- We conclude that leaves cannot be considered to be both saturated and of a homogeneous vapour isotopic composition. They are best approximated as isotopically homogeneous.

## Introduction

Leaf stable isotope compositions are an important measurement for plant scientists and agriculturalists, dating back to at least the 1960s with the first observation of isotopic enrichment of leaf water due to transpiration by Gonfiantini *et al*. (1965) and the subsequent explanation of the phenomenon through the Craig-Gordon equation as adapted to leaves (Dongmann *et al*. (1974); Farquhar *et al*. (1989)). Since then, several stable isotopes have been used to probe the structure, biochemistry and metabolism of leaves. For instance, ^2^H,^13^C, ^15^N and ^18^O, which demonstrate varying degrees of diel variability in the Australian plant species *Lupinus angustifolius* and *Eucalyptus globulus*, linked to time-varying metabolic and photosynthetic rates (Cernusak *et al*. (2002); Cernusak *et al*. (2005)). Particularly important from a leaf gas exchange perspective are the relative abundances of the isotopologues of carbon dioxide and water. The former has been used to study the photosynthesis of C_3_ and C_4_ plant species (Farquhar *et al*. (1989)) and contributed to global models of CO_2_ fluxes (Dubbert & Werner (2018)). The latter, which forms the subject of this paper, has contributed to global models of water flux (Farquhar *et al*. (1993)) and demonstrated a utility to industry, for instance in its use as a proxy for stomatal conductance to explain increased wheat crop yields with genetic selection (Barbour *et al*. (2000)).

Many modern methods of measuring isotopologues of transpired water *in planta* use an open gas exchange system with a chamber to enclose a leaf, so the isotopic composition of water vapour entering and exiting the leaf chamber can be assessed (Cernusak *et al*. (2016)). These methods have several strengths, in particular the capacity for non-destructive, in vivo and time-dependent measurements of leaves. However, chamber methods also have two common limitations: they are often predicated on the assumption of leaf saturation (Gaastra (1959); Cernusak *et al*. (2018); Rockwell *et al*. (2022)) and, due to the mixing of the air within the leaf chamber, typically report only a single value for the transpired vapour composition for the entire enclosed leaf area. If water vapour within a leaf is not isotopically homogeneous, this second limitation means that chamber measurements only report a bulk isotopic composition, which might obscure more complicated leaf behaviour.

These limitations are inconsequential if two assumptions hold: the air within a leaf is saturated with water vapour, and leaf water vapour is isotopically homogeneous (for the leaf volume enclosed in a chamber). However, to our knowledge, there is no direct evidence of the concurrent occurrence of saturation and isotopic homogeneity, nor the impact of possible unsaturation in the substomatal cavity on where the vapour in the leaf becomes inhomogeneous. Subsequently, we consider in this paper the validity of concurrently assuming these two predicates. That is, whether it is valid to assume that leaves are saturated with water vapour at a low vapour pressure deficit (1 kPa), and that this water vapour is of a homogeneous isotopic composition.

The first assumption, that air within a leaf is saturated with water vapour, has been well-studied historically, with mixed results found (Cernusak *et al*. (2024)). Some studies have shown support for unsaturation (Jarvis & Slatyer (1970); Ward & Bunce (1986); Cernusak *et al*. (2018)), and others have shown no evidence (Farquhar & Raschke (1978); Sharkey *et al*. (1982)). However, in more recent times, with Wong *et al*. (2022) reporting relative humidities in intercellular air spaces of 90% (*Gossypium hirsutum)* and 80% (*Helianthus annuus*), the argument for unsaturation is gaining increased traction, at least for the case of leaves in conditions of high vapour pressure deficit.

There has been less historical interest in the second assumption, isotopic homogeneity in the vapour phase. Certainly, in the case of tightly closed stomata with no transpiration, diffusive mixing would be expected to eventually induce isotopic homogeneity, but the issue is more complicated when a net flow is present. Although isotopic variation within water in the liquid phase has been widely reported (Farquhar & Lloyd (1993), Farquhar & Gan (2003), Cuntz *et al*. (2007), Ogée *et al*. (2007)), it is not immediately clear if such effect is reflected in the isotopic composition of the vapour phase. Restricting our consideration to the possibility of isotopic inhomogeneity between the abaxial (lower) and adaxial (upper) surfaces of a leaf (which we can directly measure), it is intuitive to think that owing to the small distances between the surfaces, the internal air space resistance, and thus isotopic fractionation, can be neglected. However, this is also uncertain with the necessity of non-negligible internal air space and hydraulic mesophyll cell membrane resistances as required for unsaturation to occur (Wong et al. (2022)).

The study detailed in this paper began by using the leaf gas exchange apparatus of Márquez *et al*. (2021) coupled with a water vapour isotope laser analyser. This setup allowed us to independently measure the relative abundance of transpired leaf water isotopologues from the abaxial and adaxial surfaces. When using this setup, an unexpected result was obtained: when the estimated vapour isotopic composition in the leaf was computed using the Craig-Gordon equation and the assumption of saturation, different results were found for the abaxial and adaxial sides of the leaf. This result was notable for seemingly indicating an isotopic inhomogeneity in the vapour within a leaf. As the Craig-Gordon model is well-tested and founded on well-established physical mechanisms, the two most likely explanations were that either the vapour was isotopically inhomogeneous or that the internal air space within the leaf was unsaturated. The result subsequently indicated that the two previously discussed assumptions of leaf isotope gas exchange measurements could not hold concurrently.

To investigate this further, we devised an analysis with two related hypotheses and applied it to measurements of the ^18^O composition of transpired water of five-week-old *Gossypium hirsutum* plants. The first hypothesis was that if we assumed saturation, then computing the isotopic composition of the evaporative site of water vapour with the Craig-Gordon model would yield a different isotopic composition for the abaxial and adaxial sides of the leaf; i.e. that assuming saturation leads to the inference of an isotopic inhomogeneity. The second hypothesis was that if we assumed isotopically homogenous water vapour, then computing the humidity of the substomatal cavities of a leaf with the Craig-Gordon model would yield a different result for the abaxial and adaxial surfaces; i.e., that assuming isotopic homogeneity leads to the inference of unsaturation. Support was found for both hypotheses, indicating that the two commonly held assumptions cannot hold concurrently.

Finally, we turned to pre-existing data from Wong *et al*. (2022) to distinguish between the two likely explanations, namely, to determine whether our results were predominantly explained by our leaves being isotopically inhomogeneous, or unsaturated. From an analysis of these data with our newly devised analysis method, we found evidence that a difference in the degree of unsaturation between the adaxial and abaxial substomatal cavities was the predominant contributor to our results.

## Theory

### Glossary of Standard Notation

### Assumed Saturation Method

To determine whether assuming saturation leads to the inference of isotopic inhomogeneity, we consider the Craig-Gordon model as adapted to leaves (Craig & Gordon (1965); Dongmann *et al*. (1974); Farquhar et al. (1989); Farquhar et al. (2021)). This model is encapsulated by the equations of flux for major (equation 1) and minor (equation 2) water isotopologues, the latter equation termed the Craig-Gordon equation:

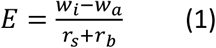

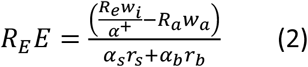

where the full list of symbols used here (and elsewhere in the paper) is given in Table 1.

**Table 1:** List of symbols used in the text, how their values are computed/assigned and whether these evaluations require the assumption of saturation or isotopic homogeneity.

| Symbol | Meaning | Assumption required to evaluate | Assigned Value / Means of Computation |
| --- | --- | --- | --- |
| $a$ | 2<sup>nd</sup> order quadratic expansion term | No | Discussed in the theory section |
| $\alpha_b$ | $^{18}\text{O}/^{16}\text{O}$ fractionation of water due to boundary layer resistance | No | $\alpha_b = 1.019$<br>(Gonfiantini *et al.* (2018)) |
| $\alpha^+$ | $^{18}\text{O}/^{16}\text{O}$ fractionation of water due to equilibrium isotope effect | No | As per Bottinga & Craig (1969) |
| $\alpha_s$ | $^{18}\text{O}/^{16}\text{O}$ fractionation of water due to stomatal resistance | No | $\alpha_s = 1.0285$<br>(Gonfiantini *et al.* (2018)) |
| $b$ | 1<sup>st</sup> order quadratic expansion term | No | Discussed in the theory section |
| $c$ | 0<sup>th</sup> order quadratic expansion term | No | Discussed in the theory section |
| $E$ | Transpiration rate per unit area | No | As per von Caemmerer & Farquhar (1981) |
| $k$ | Proportionality factor | No | Discussed in the theory section |
| $R_a$ | Relative isotopic composition ($^{18}\text{O}/^{16}\text{O}$) of water vapour in chamber | No | Directly measured |
| $R_E$ | Relative isotopic composition ($^{18}\text{O}/^{16}\text{O}$) of transpired water | No | As per Farquhar *et al.* (2021) |
| $R_e$ | Relative isotopic composition ($^{18}\text{O}/^{16}\text{O}$) of evaporative site | Yes | Discussed in the theory section |
| $R_x$ | Relative isotopic composition ($^{18}\text{O}/^{16}\text{O}$) of xylem water | - | Value not assigned /computed (used only in Figure 9) |
| $r_b$ | Boundary layer resistance to water vapour | No | Empirically determined |
| $r_{ias}$ | Internal air space resistance to water vapour | - | Value not assigned /computed (used only in Figure 9) |
| $r_s$ | Stomatal resistance to water vapour | Yes | Discussed in the theory section |
| Subscript ab | Pertains to the abaxial (lower) surface of a leaf | N/A | N/A |
| Subscript ad | Pertains to the adaxial (upper) surface of a leaf | N/A | N/A |
| Subscript ext | Pertains to measurements made by a method external to this paper | N/A | N/A |
| $w_a$ | Water vapour concentration in the air inside the chamber | No | Directly measured |
| $w_i$ | Water vapour concentration in the substomatal cavity | Yes | Discussed in the theory section |
| $w_s$ | Water vapour concentration in the air at the leaf surface | - | Value not assigned /computed (used only in Figure 9) |
| $w_{sat}$ | Water vapour concentration at 100% relative humidity | No | As per equation 10 of Bolton (1980) |

These equations are widely used in leaf isotope gas exchange measurements to compute the isotopic composition of liquid water at the evaporative sites within a leaf, *R*_*e*_, assuming that the water vapour concentration in a leaf’s substomatal cavity, *w*_*i*_, is saturated with water vapour at leaf temperature, that is: *w*_i_ = *w*_sat_ (see for example Farquhar *et al*. (2007)). In this common usage, the entire leaf is considered as a single system, meaning, for example, that transpiration is considered as the sum of all contributions from the abaxial and adaxial leaf surfaces. In such a treatment only a single value for the isotopic composition of the leaf’s evaporative sites is ascertained.

However, despite this common usage for combined surfaces, these equations can also be used to consider each surface of the leaf independently (indeed, it was from such a consideration that they were derived). Such an argument can be rationalised by viewing the equations as a mathematical encoding of known physical principles. In particular, they are recognisable as simple statements of Fick’s first law (Fick 1855), with the Craig-Gordon equation also incorporating a quantification of the kinetic and equilibrium isotope effects (Craig *et al*. (1963); Gonfiantini *et al*. (2018)). Notably, the validity of these principles is not based on the geometry of a leaf, so the equations are also valid when considering each surface of the leaf independently.

Subsequently, we use these two equations to consider each surface of the leaf independently. Specifically, we rearrange equation 1 to solve for the stomatal resistance of a single side of the leaf as follows:

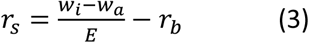

And then use this result with the Craig-Gordon equation to solve for the isotopic composition of the evaporative site, *R*_*e*_ for each surface:

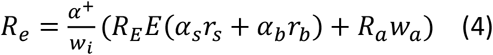

For each surface, we assume that *w*_*i*_ = *w*_*sat*_ within each surface of the leaf.

In a hypothetical experiment, all terms on the right-hand side of equation 4 except for the fractionation factors (*α*_*s*_, *α*_*b*_ and *α*^+^) and *w*_*i*_ could differ between the two surfaces of the leaf. However, if the leaf is isotopically homogeneous and saturated, then these differences would exactly cancel to produce the same *R*_*e*_ for both surfaces. Subsequently, if a statistically significant difference is found between the values of *R*_*e*_ for both surfaces of the leaf, then we find that our leaf is not isotopically homogeneous if it is saturated, which contradicts the concurrent assumption of both isotopic homogeneity and saturation.

### Assumed Isotopic Homogeneity Method

To determine whether assuming isotopic homogeneity leads to the inference of unsaturation, we again begin with the Craig-Gordon model as adapted to leaves. As before we note that each side of the leaf can be treated independently in the equations of water flux (equations 1 and 2) and, by the same argument, note that these equations are equally applicable to the case of a non-saturated *w*_i_.

Subsequently, we establish the validity of equations 1 and 2 and hence their rearrangements, equations 3 and 4, for the case when *w*_*i*_ < *w*_sat_.

Now, in the assumed saturation method, we had a value for *w*_*i*_ (*w*_sat_) which allowed us to solve equation 3. However, when assuming isotopic homogeneity, we instead need to solve for *w*_*i*_. Thus, rather than solving equation 3 immediately, we will combine it with equation 4 via substitution and continue our derivation:

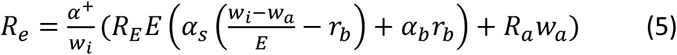

Following a rearrangement of this equation for 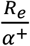 and simplification, we arrive at the result:

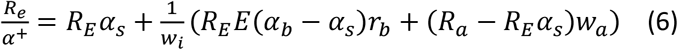

For reasons of brevity, we will define *k* = *R*_*E*_*E*(*α*_*b*_ − *α*_*s*_)*r*_*b*_ + (*R*_*a*_ − *R*_*E*_*α*_*s*_)*w*_*a*_ and substitute it into the above expression giving the result:

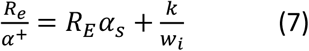

Now to proceed with the development of our method, we recognise that *R*_*e*_ is the isotopic composition of liquid water at the evaporative site, and *α*^+^ is the coefficient that relates this isotopic composition to water vapour at the liquid-air interface of such sites. Subsequently, the term 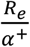 is recognisable as the isotopic composition of water vapour at the liquid-air interface.

At this point we invoke our assumption of isotopic homogeneity, noting that all water vapour within the leaf must take the same isotopic composition. In particular then, this 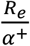 term must be the same for values from both sides of the leaf so we can equate the right-hand side of equation 7 considering terms restricted to a particular side. Adopting the convention of subscripts *ab* and *ad* to distinguish between values from the abaxial and adaxial surfaces, respectively, we arrive at:

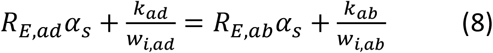

Now we can rearrange this equation for *w*_*i,ad*_, arriving at the following result:

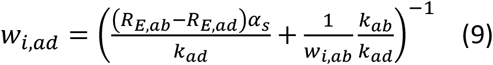

Equation 9 is a parametric equation relating the two variables of interest, *w*_*i,ad*_ and *w*_*i,ab*_. This means that, in particular, it does not return a given value for the water vapour concentration in the substomatal cavity when computed for each surface, but an infinite family of possible pairs of adaxial and abaxial *w*_i_. It bears noting that this different behaviour of the solution as compared to the assumed saturation method arises not from a degeneracy in the Craig-Gordon model, but is a reflection of the fact that assuming isotopic homogeneity is an inherently weaker assumption than assuming saturation. Specifically, assuming saturation assigns a value (*w*_sat_) to both *w*_*i,ab*_ and *w*_*i,ad*_ (2 independent constraints), whereas assuming isotopic homogeneity only assigns a value to *R*_*e,ab*_, namely *R*_*e,ad*_ (1 independent constraint). This asymmetry means that our assumed isotopic homogeneity method should have an additional parameter, exactly as observed.

#### The Maximal Humidity Constraint

To rectify the problem of an unconstrained parameter and derive a single solution from equation 9, it is necessary to impose an additional constraint. Many candidate constraints could be envisioned, but for the purpose of testing whether assuming isotopic homogeneity leads to the inference of unsaturation, we consider a maximal humidity constraint. This entails choosing the solution of equation 9 with the greatest combined *w*_i_ of both sides of the leaf without exceeding *w*_sat_ in the substomatal cavity of either surface (i.e., *w*_i_/*w*_sat_ ≤ 1). Importantly for our purposes, imposing such a maximal humidity constraint means that if there exists a solution with *w*_i_ = *w*_sat_ when assuming isotopic homogeneity, i.e. a solution not indicative of unsaturation, then it will be selected.

To draw a statistical conclusion on the existence of unsaturation with this constraint is then a simple task. We consider a null hypothesis that *w*_i_=*w*_sat_ on either side of the leaf and that a given measurement would have some symmetric uncertainty. As an alternative hypothesis, we consider *w*_i_<*w*_sat_. Given that we constrain our *w*_i_ to be less than or equal to *w*_sat_, then according to the null hypothesis we have, for a given measurement, a probability *p* = 0.5 of measuring *w*_i_=*w*_sat_, and a probability 1 − *p* = 0.5 of measuring *w*_i_<*w*_sat_. Thus, over several measurements, the number of measurements of *w*_i_=*w*_sat_ under the null hypothesis is given by a binomial distribution. Determining the p-values for a given set of measurements will then allow us to conclude if there is evidence of unsaturation. This approach is subsequently used to test for the existence of unsaturation when isotopic homogeneity is assumed.

#### The External Estimate Constraint

An alternative constraint for the solution of equation 9, and one that we will use to argue for the validity of assuming isotopic homogeneity, is found by measuring bulk leaf unsaturation using an external method, for example the method of Cernusak *et al*. (2018). This external measurement can then be used to constrain the solutions of the parametric equation by imposing the condition that:

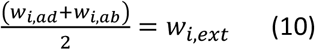

where *w*_*i,ext*_ is some measurement of substomatal cavity water concentration by an external method.

In such a case, equation 9, coupled with the above constraint gives two prospective solutions of the form:

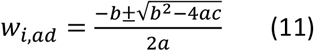

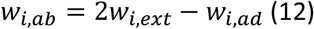

where:

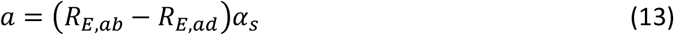

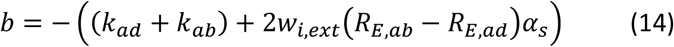

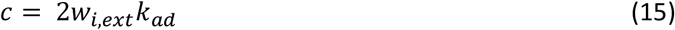

The physically realistic solution can be determined by observation (for example, identifying the solution predicting non-negative *w*_i_). The derivation of these solutions is given in Note 1 of the supporting information.

## Materials and Methods

### Plant Materials

Seeds of *Gossypium hirsutum* (cotton) were sown in late July – mid August 2022 (the austral winter) into 10 L plastic pots containing steam-sterilised potting mix and fertilised with slow-release fertiliser (Osmocote; Scotts, New South Wales). The seeds were germinated and grown in a greenhouse on the Australian National University Acton campus and thinned to one plant per pot to produce a sample of uniform plants. The plants were exposed to natural light conditions, with noon-day irradiance in the photosynthetically active range at the uppermost leaves of the plants of approximately 1200 μmol m^-2^ s^-1^. A temperature control was used to maintain a daytime air temperature of 28 ± 2°C and a night-time air temperature of 20 ± 2°C. The plants were well watered during their growth and used for measurements approximately 5 weeks after being sown.

### Gas Exchange Apparatus

A simplified schematic of the apparatus for gas exchange measurements is shown in Figure 1. A detailed description of the chamber set-up is available in Márquez *et al*. (2023b). In brief summary, compressed air was scrubbed of H_2_O and CO_2_ and supplied to two Li6800 consoles (Li-Cor, Nebraska). Using a cylinder of pure CO_2_ and a column humidifier filled with reverse osmosis filtered water (of δ^18^O value approximately -5‰), the Li-Cor consoles reintroduced CO_2_ and H_2_O to the air in controlled concentrations. This air was then directed through a Li6800-01A head unit (Li-Cor, Nebraska) to chambers clamped onto the abaxial and adaxial surfaces of a mature cotton leaf. The air in the chambers exchanged gases with the photosynthesising leaf and was then redirected back through the Li6800-01A head unit. A portion of the ingoing and outgoing air fed to each leaf chamber was redirected to a four-way valve leading to a Picarro cavity ringdown spectrometer (L2130-I Isotopic H_2_O; Picarro, California) and an Aerodyne dual quantum cascade laser (QCLAS-ISO; Aerodyne Research Inc, Massachusetts).

**Figure 1:**
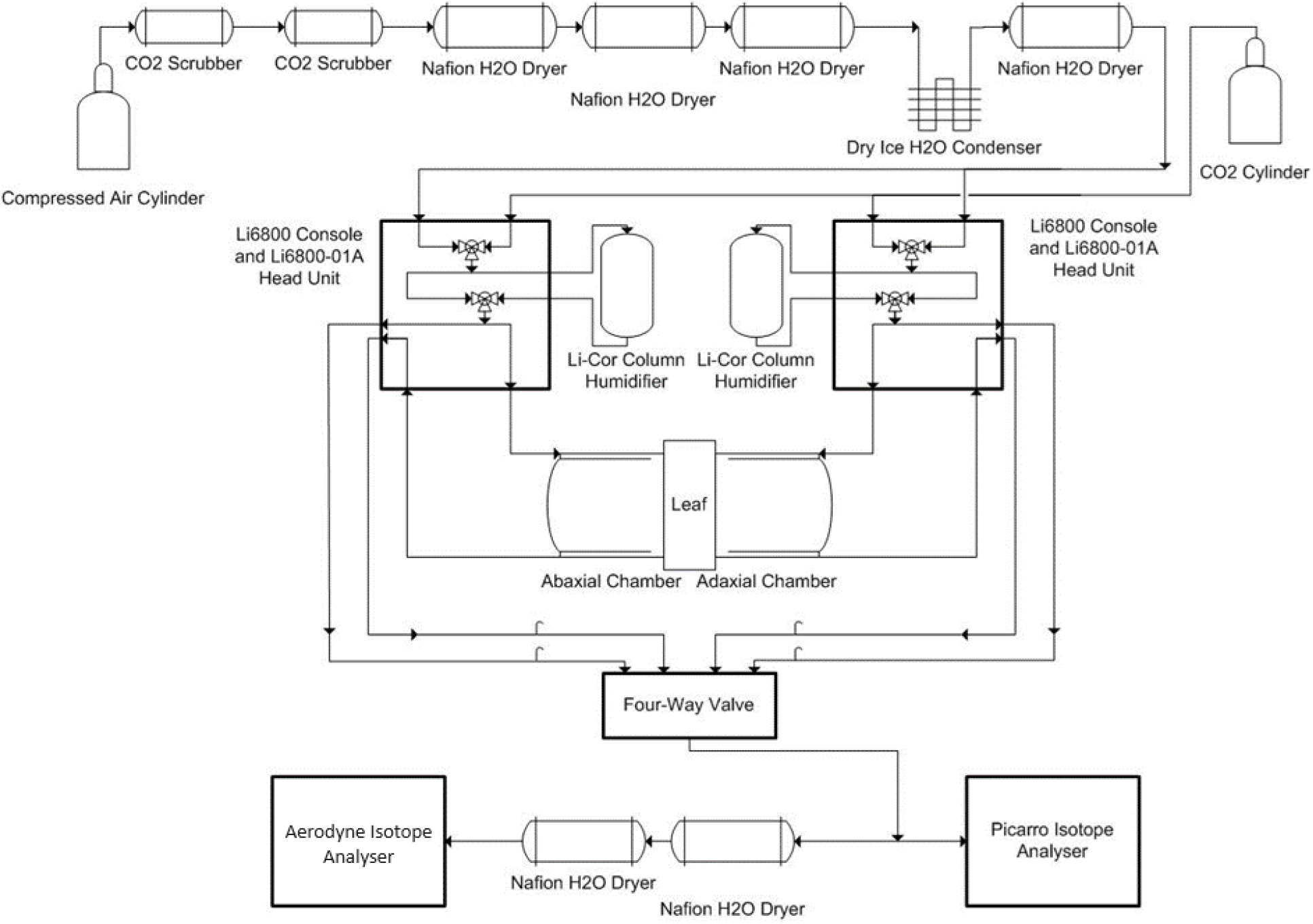
Simplified schematic of gas-exchange apparatus.

The humidity and CO_2_ content of the ingoing and outgoing air to each of the leaf chambers was measured using the infrared gas analysers in the Li6800-01A head units. The δ ^18^O-composition of the H_2_O was measured using the Picarro isotope analyser. Note that no measurements from the Aerodyne were used in the analysis of newly collected data; the instrument was included in the apparatus as it is the normal setup and to draw a greater flow of air through the four-way valve (∼260 ml min^-1^).

### Gas-Exchange Method

Gas-exchange measurements were made from the 8^th^ of September to the 11^th^ of October 2022 (the austral spring). Plants were moved the night before measurements into a lab with a temperature of 23.5°C, and 1500 μmol m^-2^ s^-1^ of photosynthetically active light (as measured with a Li-190SB-L photometer; Li-Cor, Nebraska) was provided to the top surface of the leaves by a ViparSpectra XS1000 solar lamp (ViparSpectra, California) between 06:00 and 18:00.

Measurements were performed on mature leaves of cotton plants under conditions of 1500 μmol m^-2^ s^-1^ of light intensity from the top with 96% red and 4% blue light, at an air temperature of approximately 24°C, at a chamber pressure 0.2 kPa greater than the atmosphere, at a CO_2_ concentration of 400 ppm and at a 1.0 kPa air saturation deficit (an H_2_O concentration of ∼20,000 ppm). The fans in both chambers were set to rotate at 10,000 rpm and the flow of air into each chamber was set to 500 μmol s^-1^.

The leaf was left to reach gas exchange steady state at the set conditions in the gas exchange chambers. Steady-state was considered as a < 1% change over a 16-minute period in the assimilation rate of CO_2_ and stomatal conductance to water of both the abaxial and adaxial leaf surfaces. Then, the gas exchange data from the Li6800-01A head units were continually recorded and the isotopic measurements with the Picarro isotope analyser were begun. For each sample of gas, isotopic measurements were taken as the average of a four-minute period in isotopic steady-state, considered for this study as > 95% of points of δ^18^O for H_2_O remaining within a 0.5‰ band. To take these isotopic measurements, the four-way valve (see Figure 1) was rotated to select each of the four gas portions to be sampled, with the direction of rotation and first sample considered alternating between days. For each leaf, it took up to 4 hours to achieve the required level of stability and begin taking these measurements.

In total, 27 sets of measurements of each of the four gas portions were taken across 8 distinct leaves.

## Results

### Assuming Saturation Yields Isotopic Inhomogeneity

Results are shown in Figure 2 and Table S1 for the computation of the isotopic composition of the liquid water at the evaporative sites of the abaxial and adaxial surfaces using our assumed saturation method on our newly collected data. The distribution of the data is shown by a box and whisker plot. Here and elsewhere in the paper, the whiskers extend from the first and third quartile out to 1.5 times the inter-quartile range or the most outlying data point, whichever is closer to the median. The median is shown as the central orange line, and (where present) outliers are shown by open circles.

**Figure 2:**
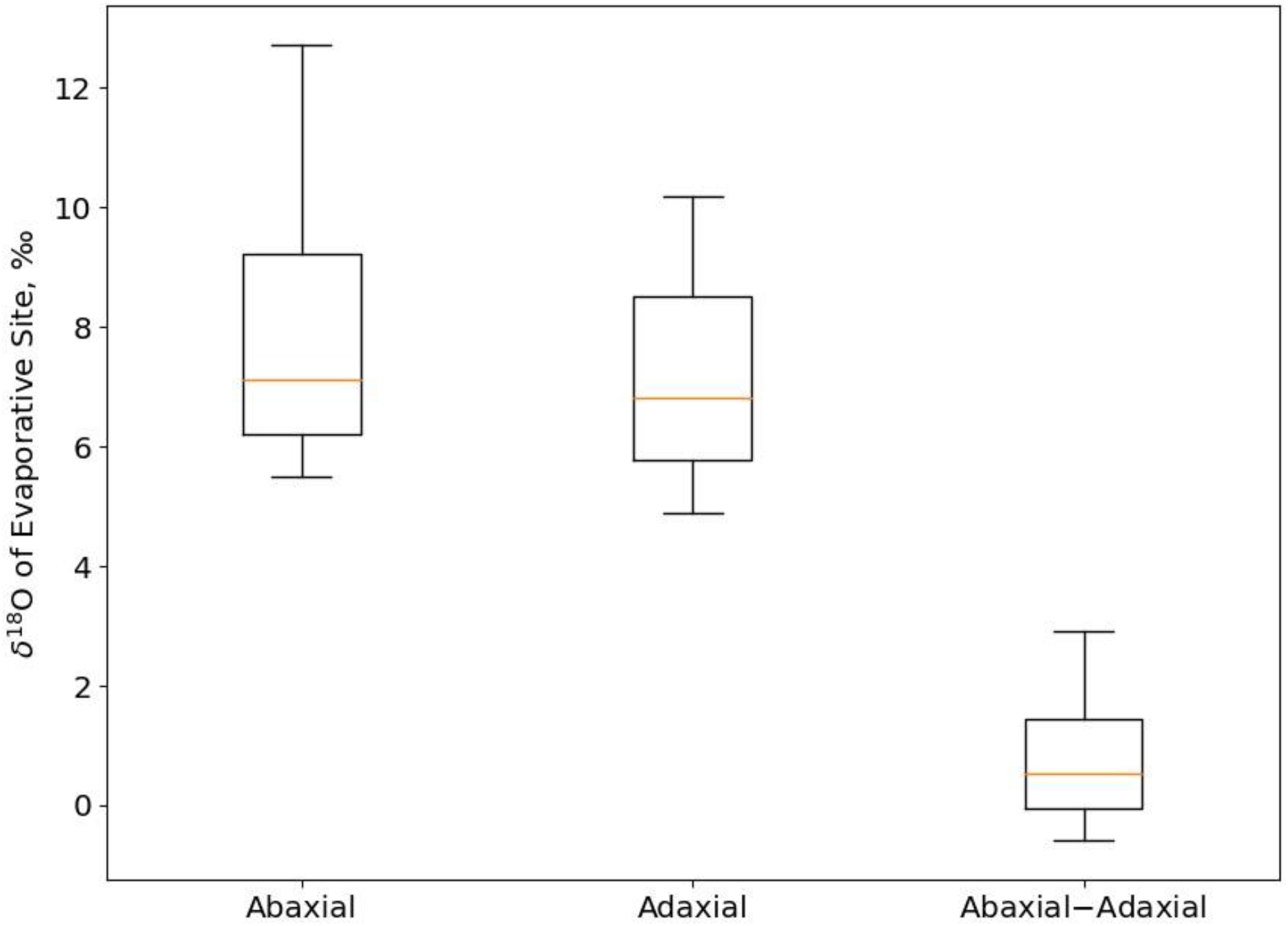
isotopic composition (*δ*^18^O) of the abaxial and adaxial evaporative sites, and their difference, for *gossypium hirsutum* at 400ppm CO_2_ and 1.0 kpa air saturation deficit when assuming that the internal air spaces of leaves are saturated. a statistically significant difference was found between the isotopic compositions of the abaxial and adaxial surfaces (p-value 0.04).

A statistically significant difference to a threshold of 95% confidence was observed between the abaxial and adaxial surfaces (P-value: 0.04) by a two-tailed paired-samples t-test. Thus, the results indicate that assuming saturation yields the calculation of an isotopic inhomogeneity within the leaf.

### Assumed Isotopic Homogeneity Yields Unsaturation

Results are shown in Figure 3 and Table S1 for the computation of the humidities in the abaxial and adaxial substomatal cavities of leaves using our assumed isotopic homogeneity method and the maximal humidity constraint on our newly collected data. Also shown are the computed stomatal resistances of the abaxial and adaxial surfaces for use in the discussion. Figure 4 and Table S2 show three examples of the parametric curves derived when not applying an additional constraint to our assumed isotopic homogeneity method (equation 9). The three leaves shown (leaves B, C and D) from the newly collected data were chosen as they have the greatest number of repeated measurements, but the other leaves present the same behaviour.

**Figure 3:**
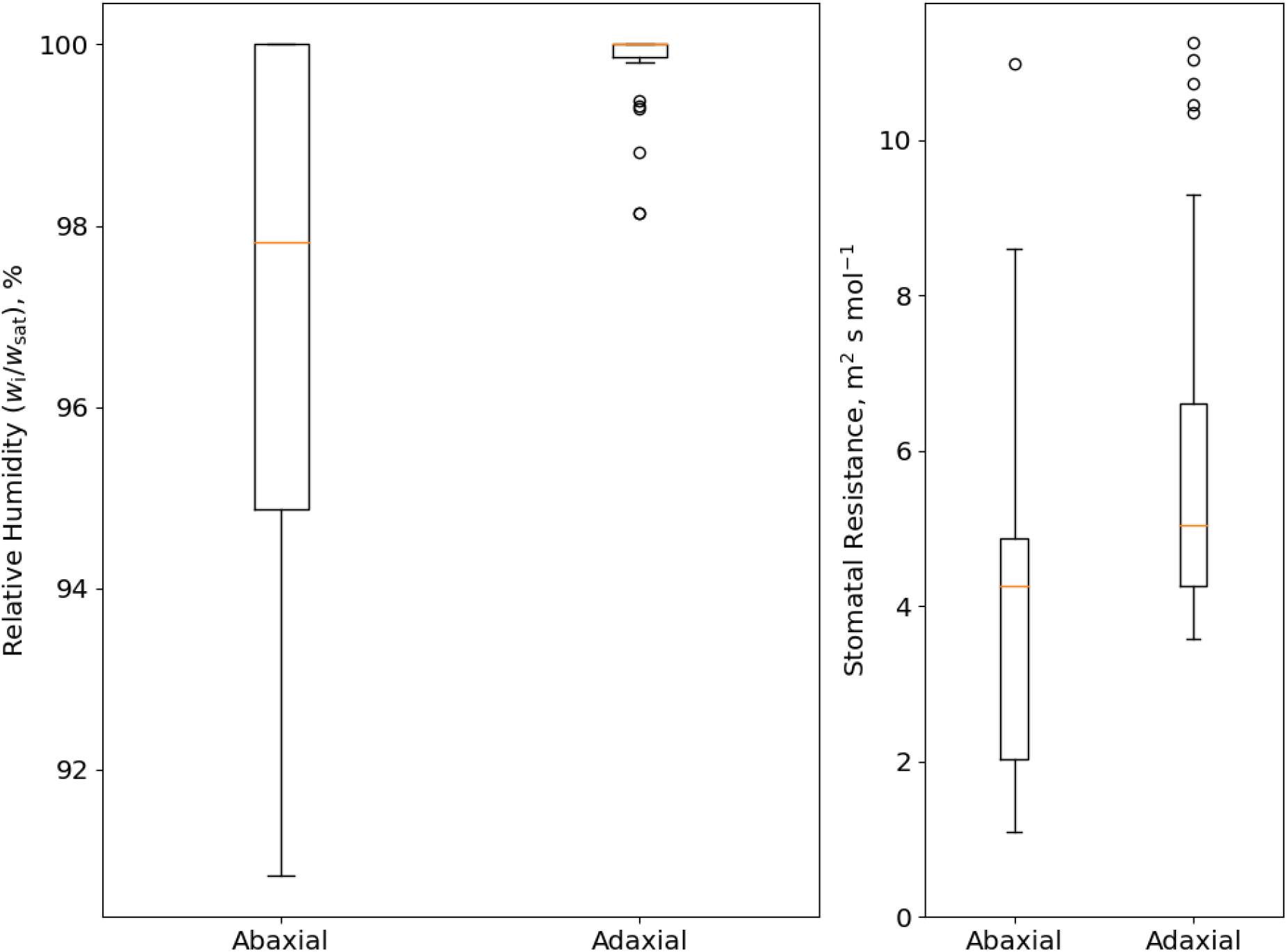
(Left) Relative humidity in the leaf substomatal cavity (*w*_i_/*w*_sat_) of the abaxial and adaxial substomatal cavities computed using the assumed isotopic homogeneity method with the maximal humidity constraint. (Right) stomatal resistance of the abaxial and adaxial surfaces computed using the same method. For both cases, data are for *Gossypium hirsutum* leaves at 400 ppm CO_2_ and 1.0 kPa air saturation deficit. A statistically significant difference between our measurements and saturation was found for the abaxial surface (P-value: 0.03). No statistically significant difference was found for the adaxial surface (P-value: 0.99).

**Figure 4:**
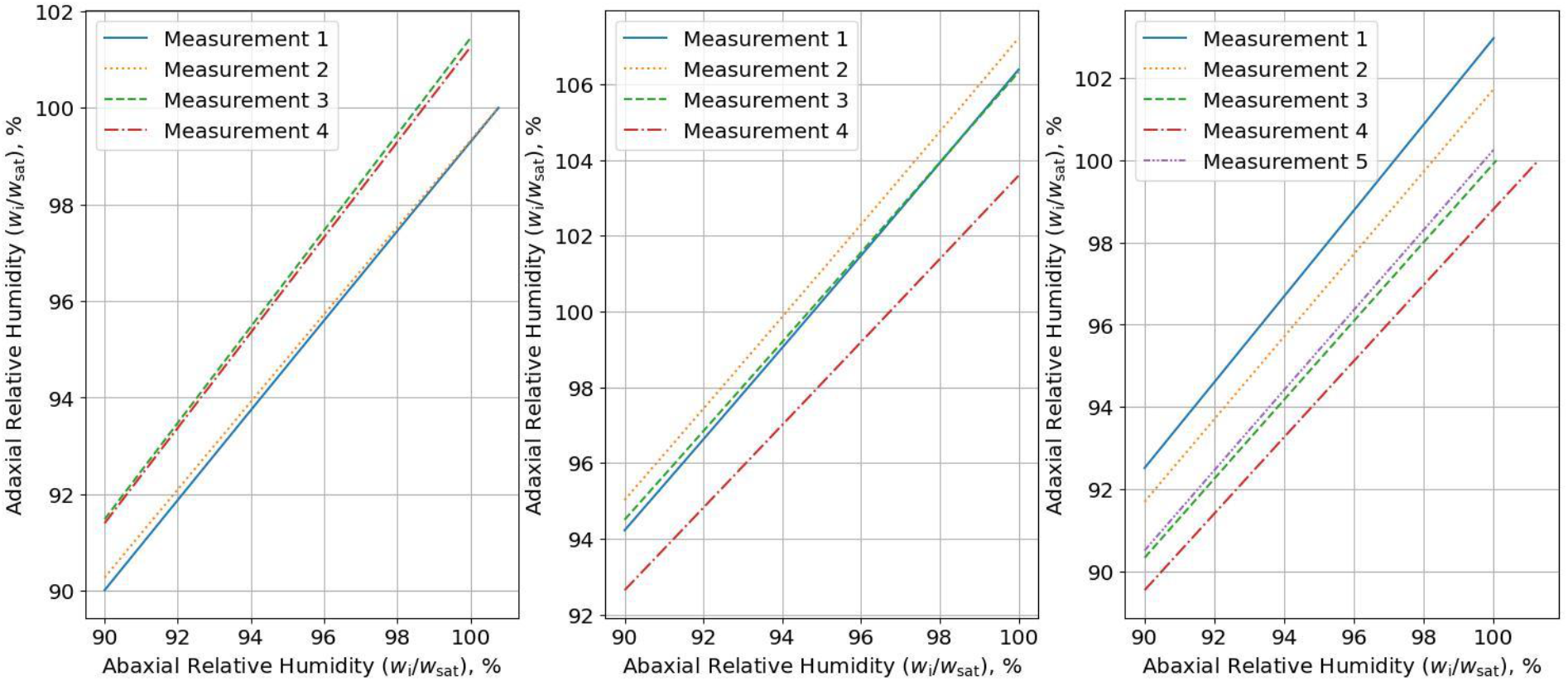
Three examples of the parametric curves generated from equation 9, constituting the assumed isotopic homogeneity method without an additional constraint. Going from left to right, leaves B, C and D of the new data for *Gossypium hirsutum* leaves at 400 ppm CO_2_ and 1.0 kPa air saturation deficit are shown. Note the positive correlation between the relative humidity of the abaxial and adaxial substomatal cavities (*w*_i_/*w*_sat_) and the greater unsaturation of the abaxial surface relative to the adaxial.

Applying the hypothesis test detailed in the theory section to the data shown in Figure 3, we find a statistically significant difference (P-value: 0.03) between the number of abaxial leaf measurements at *w*_i_=*w*_sat_, and what would be expected were the leaves saturated. Thus, the results provide evidence that assuming isotopic homogeneity leads to the inference of unsaturation. We find no statistically significant difference (P-value: 0.99) between the number of adaxial leaf measurements found at *w*_i_=*w*_sat_ and the expectations of a saturated leaf. This is consistent with Figure 4, which shows our parametric equation predicts a *w*_i_ closer to *w*_sat_ for the adaxial surface.

### Evidence in Favour of Assuming Isotopic Homogeneity

To consider whether assuming saturation or isotopic homogeneity is preferable, we examined a pre-existing dataset on *Gossypium hirsutum* from Wong *et al. (*2022) for the purpose of having an independent measure of saturation. In this dataset, the two leaf sides were again measured independently, and the *δ*^18^O of both the water vapour and CO_2_ entering and exiting each cuvette were determined.

In Figure 5 and Table S3 we show the difference between computed isotopic compositions of liquid water at the evaporative sites of the abaxial and adaxial surfaces using the assumed saturation method against the mean *w*_i_ (combining abaxial and adaxial measurements) reported by Wong *et al*. (2022) using the Cernusak *et al*. (2018) method. Notably there is a statistically significant monotonic correlation to a threshold of 95 % confidence (P-value: 0.02) by a Kendall’s Tau correlation test against the null hypothesis of no correlation. More specifically, the difference in isotopic composition between the two surfaces declines with increasing saturation (i.e., *w*_i_ approach *w*_sat_). Taking a simplistic linear relation, this reaches a value of ∼0.8‰ at *w*_i_ = *w*_sat_ (R^2^=0.29).

**Figure 5:**
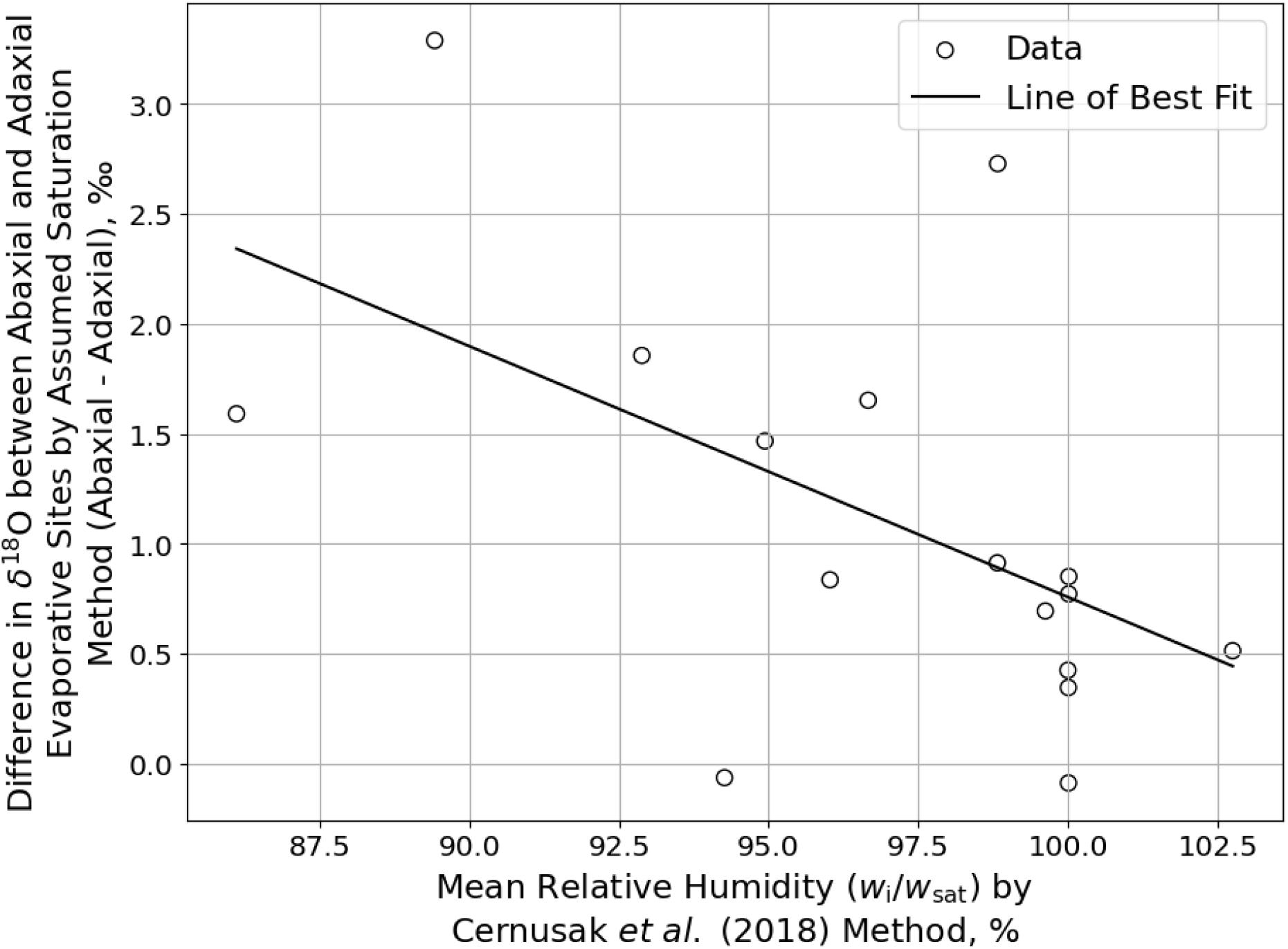
Difference in isotopic composition (*δ*^18^O) between the abaxial and adaxial evaporative sites of *Gossypium hirsutum* leaves as determined by the assumed saturation method versus the mean *w*_i_/*w*_sat_ of the leaf as determined by the Cernusak *et al*. (2018) method. By Kendall’s Tau correlation test, there is a statistically significant monotonic correlation to a threshold of 95% confidence (P-value: 0.02) against the null hypothesis of no correlation. The linear fit has R^2^=0.29. Data are sourced from Wong *et al*. (2022).

In Figure 6 and Table S3 we show the required *w*_i_/*w*_sat_ of the abaxial and adaxial substomatal cavities to match the results of Wong *et al*. (2022) and be consistent with isotopic homogeneity; our assumed isotopic homogeneity method with the external estimate constraint. The *w*_i_ at the adaxial substomatal cavity is notably maintained at approximately *w*_sat_, even when the average humidity (*w*_i_/*w*_sat_) reduces to 90%. By contrast, the *w*_i_/*w*_sat_ of the abaxial substomatal cavity steadily decreases with overall leaf unsaturation. The predicted *w*_i_/*w*_sat_ of the adaxial substomatal cavity exceeds unity for some measurements, which is commented on in the discussion. Also pictured is the stomatal resistance computed by the assumed isotopic homogeneity method for use in the discussion.

**Figure 6:**
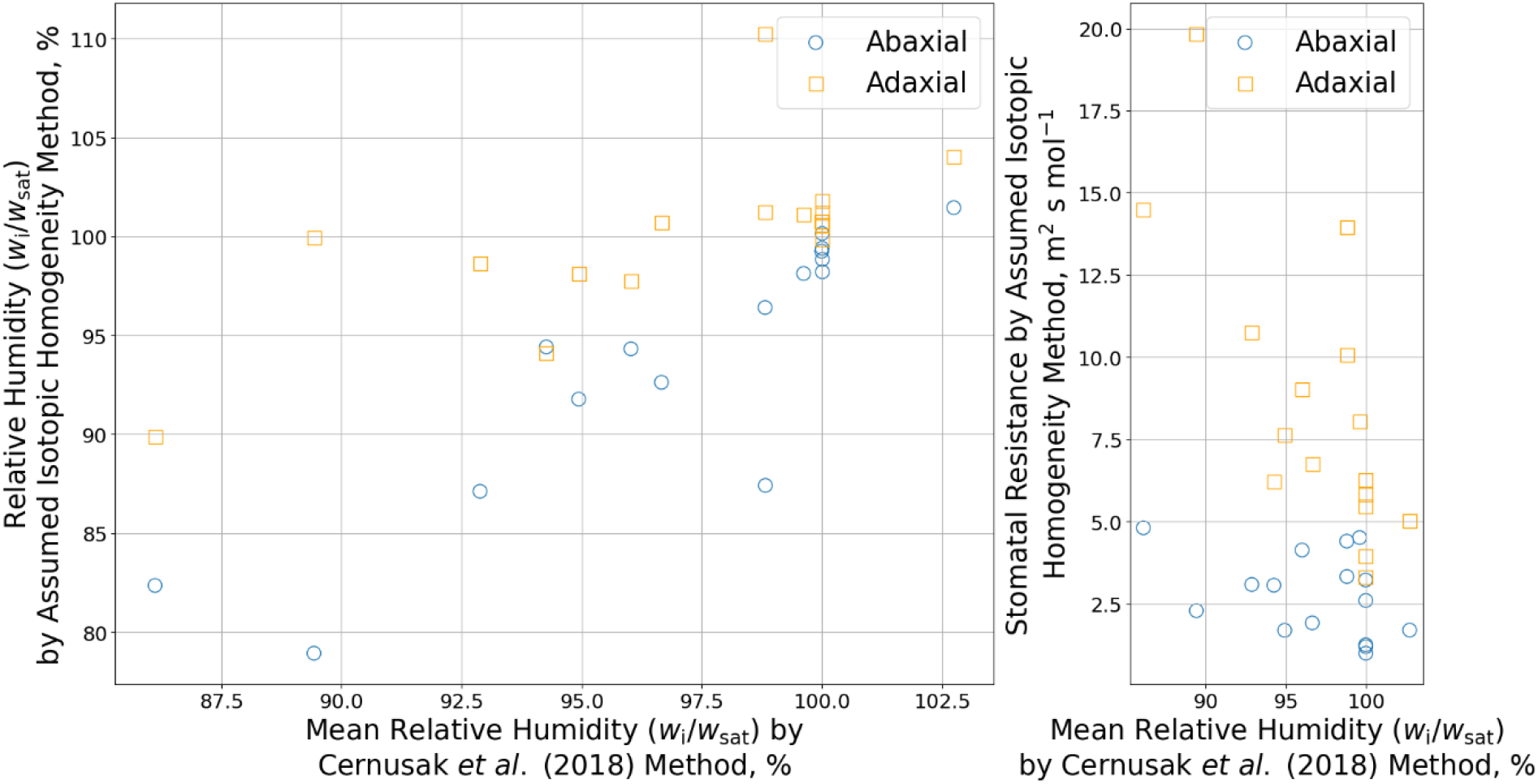
(Left) computed relative humidity of abaxial and adaxial substomatal cavities (*w*_i_/*w*_sat_) using the assumed isotopic homogeneity method with the reported *w*_i_ from the Cernusak *et al*. (2018) method as a constraint and (right) stomatal resistances from the same calculations. Note the greater *w*_i_/*w*_sat_ and stomatal resistance of the adaxial surface. Data are sourced from Wong *et al*. (2022).

Results are shown in Figure 7 and Table S3 for computations of the *w*_i_ at the abaxial and adaxial substomatal cavities using the assumed isotopic homogeneity method and the maximal humidity constraint against the reported values from Wong *et al*. (2022) using the Cernusak *et al*. (2018) method. No statistically significant difference between the two sets of values was found by a two-tailed paired-samples t-test (P-value: 0.76). This suggests that the two measurements of unsaturation are equivalent within uncertainties.

**Figure 7:**
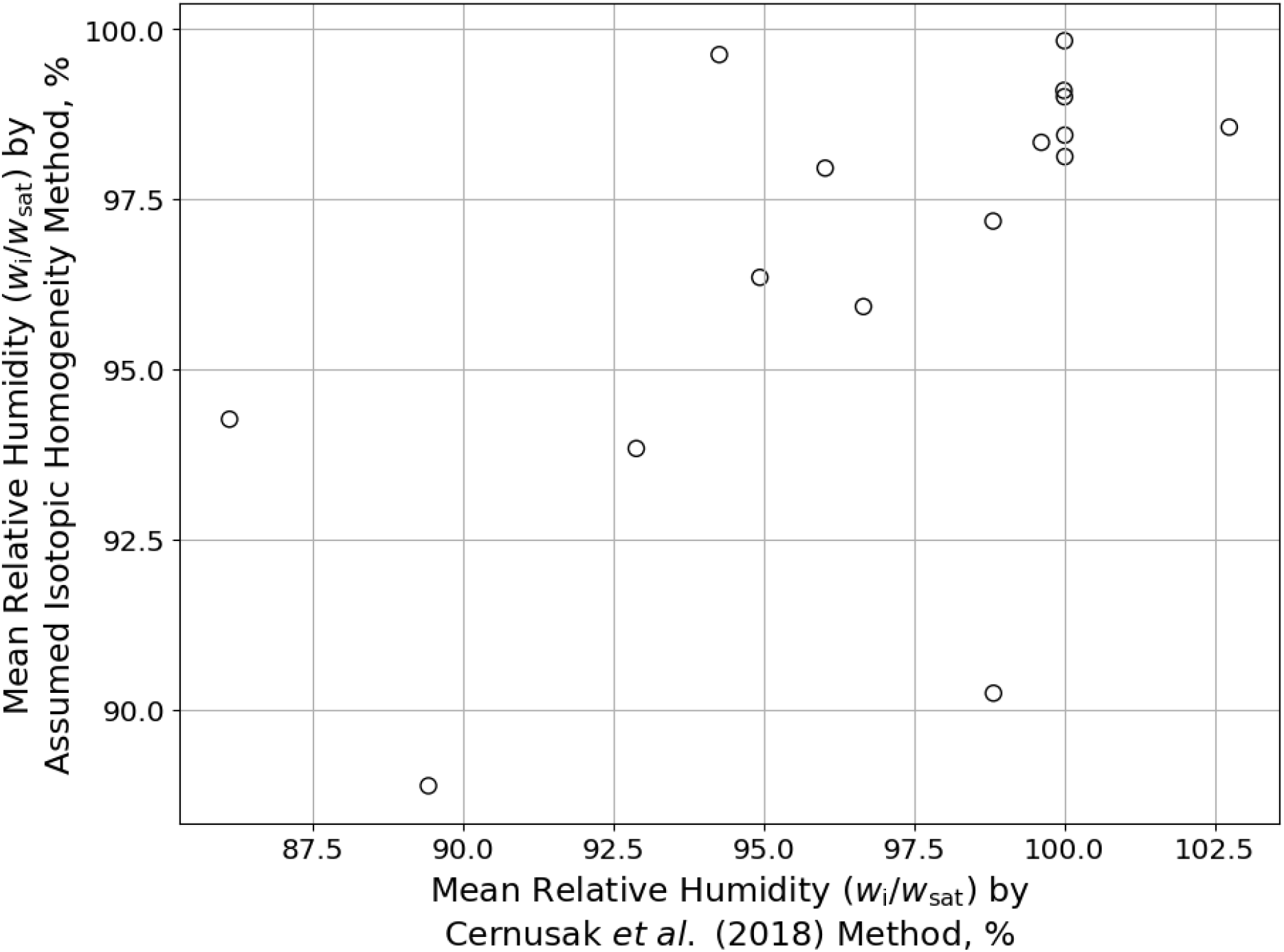
Mean relative humidity of substomatal cavities (*w*_i_/*w*_sat_) using the assumed isotopic homogeneity method with the maximal humidity constraint versus the percentage saturation from the Cernusak *et al*. (2018) method. There was no statistically significant difference between the two measures to 95% confidence (P-value: 0.76) by a two-tailed paired-samples t-test. Data are sourced from Wong *et al*. (2022).

Finally, in Figure 8 and Table S1 we show the isotopic composition of the evaporative site as determined by the assumed isotopic homogeneity method with maximal humidity constraint against the isotopic compositions from the assumed saturation method for the newly collected data. The average difference is 0.13‰ and 0.9‰ from the adaxial and abaxial evaporative sites’ measurements when assuming saturation, respectively. For reference, the standard deviation in the mean of the abaxial and adaxial evaporative sites’ isotopic composition is 1.9‰.

**Figure 8:**
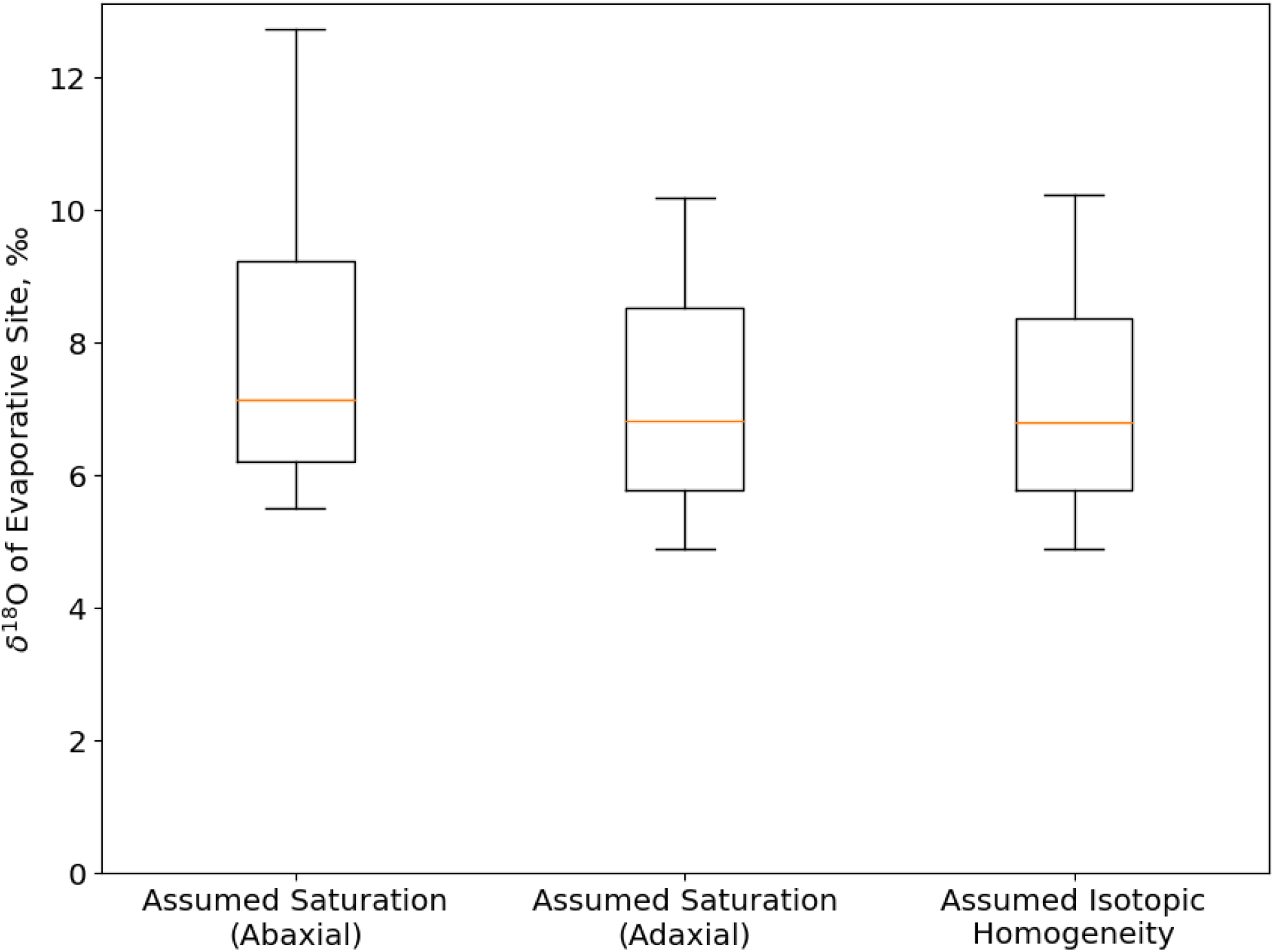
Isotopic composition (*δ*^18^O) of the evaporative sites for the abaxial and adaxial surfaces of a leaf, as computed using the assumed saturation method, and the isotopic composition of the evaporative site for the whole leaf as computed using the assumed isotopic homogeneity method with the maximal humidity constraint, for the newly collected data.

## Discussion

We found that at low air saturation deficit (1.0 kPa) assuming saturation yields the computation of an isotopic inhomogeneity within the leaf, and conversely that assuming isotopic homogeneity yields the computation of unsaturation within the leaf. Therefore, the two assumptions of isotopic leaf gas exchange measurement, namely a homogeneous isotopic composition of water vapour within the leaf and saturation of the substomatal cavities, are not self-consistent at the relatively low air saturation deficit conditions we considered in this paper. This suggests that the two assumptions cannot be satisfied concurrently.

Subsequently, we provide evidence for the existence of an inhomogeneity in either humidity or isotopic composition within the leaf, more specifically, between the adaxial and abaxial substomatal cavities. Now importantly, such an inhomogeneity can only exist if there is some non-negligible internal air space resistance within the leaf. Such a proposal is not new, with a measurement of such a resistance made by Jarvis and Slatyer (1970), Wong *et al*. (2022), and others. However, it is important to note that this resistance, along with the hydraulic resistance in the membrane cell, would inevitably lead to inhomogeneous humidity distribution within the leaf as reported by Wong *et al*. (2022). This, in turn, is likely to result in some degree of isotopic inhomogeneity as well. Thus, our results suggest that a leaf (under arbitrary conditions) is likely neither saturated nor entirely isotopically homogeneous. Subsequently we find evidence that in addition to the two assumptions not holding concurrently, they cannot be expected to hold in isolation.

Ideally, we would possess a predictive model of the internal resistances through a transect of the leaf. This could then be used to determine how unsaturation and isotopic fractionation are induced between the abaxial and adaxial surfaces (see for instance Márquez *et al*. (2023a) for an example implementation in CO_2_ measurements). However, to our knowledge no such predictive model exists in the literature. In the absence of a predictive model, it is desirable to deduce which of the two assumptions it is preferable to retain and then whether its retention is a reasonable approximation. The existence of such an approximation would be valuable as a step towards the development of such a predictive model, would allow us to comment on the spatial distribution of inhomogeneities within a leaf and, from a practical standpoint, would provide the necessary constraints to make a single-valued measurement when considering both leaf surfaces independently. Here, we lay out an argument for the usefulness of retaining the assumption of isotopic homogeneity.

### Leaves are Reasonably Approximated as Isotopically Homogeneous

This argument consists of three phases. First, that the literature supports assuming isotopic homogeneity over saturation. Second, that unsaturation explains most of the observed isotopic inhomogeneity in this paper. Third, that assuming isotopic homogeneity can reproduce results in the literature on unsaturation and (qualitatively) isotopic composition. Thus, we conclude that assuming isotopic homogeneity is a reasonable simplification.

Firstly, to consider the issue of whether it is better to assume saturation or isotopic homogeneity, we note the existence of strong empirical evidence of unsaturation in the absence of isotopic measurements (see Cernusak et al. (2024)). By contrast, to our knowledge there are no reports of a significant isotopic inhomogeneity of water vapour within a leaf. Thus, we claim that of the two common assumptions, it is preferable to retain isotopic homogeneity.

Secondly, we consider whether the occurrence of unsaturation can explain the measurement of isotopic inhomogeneity. If wrongly assuming saturation is the explanation, then any measured isotopic difference when assuming saturation must arise from a difference between the assumed and true humidities (*w*_i_/*w*_sat_). We find that a mean humidity of ∼97% would be required to explain the observed isotopic inhomogeneity for the newly collected data (Figure 3 and Table S1), which is within the range of likely substomatal cavity humidities. More stringently testing this notion with the data of Wong *et al*. (2022), the explanation still holds; from the results of Figure 5, we observe that when substomatal cavity humidities increase, as computed by the method of Cernusak *et al*. (2018), the difference in isotopic composition between the abaxial and adaxial substomatal cavities (assuming *w*_i_=*w*_sat_) decreases. This is precisely the behaviour that would be expected if the measured difference was caused by incorrectly assuming saturation. We can also consider the spatial distribution of humidities that would be required to explain such a measured difference between the abaxial and adaxial substomatal cavities (Figure 6). We find that *w*_i_/*w*_sat_ of the adaxial substomatal cavity remains at a value unity, while the reduction in *w*_i_/*w*_sat_ measured by the Cernusak *et al*. (2018) method is found at the abaxial surface. Such a spatial distribution in addition to the 2-4 times greater stomatal resistances observed on the adaxial leaf surfaces than those of the abaxial surface suggests a negative relation between substomatal *w*_i_/*w*_sat_ drop and stomatal resistance. This makes intuitive sense; if the humidity were the same for both surfaces, water vapour would transpire much faster from the lower stomatal resistance (abaxial) surface and subsequently evaporation from internal water sources would be less able to meet the transpirative demand. This would drive a *w*_i_/*w*_sat_ drop on the abaxial surface, as observed. Thus, wrongfully assuming saturation appears to result in the inference of an isotopic inhomogeneity.

As an aside, it does bear comment that in approximately half of the measurements considered in Figure 6, we observed that the adaxial surface’s *w*_i_/*w*_sat_ needed to be in ∼1-2% excess of unity. Such supersaturation is unrealistic, but it is an expected outcome owing to the assumptions and uncertainties in the combined measurements performed for this study. We made the naïve assumption that the abaxial and adaxial surfaces contribute equally to the total leaf water vapour saturation; this may not be entirely correct, and indeed a preferential weighting to the adaxial surface corrects this issue. In addition, the adaxial surface has less transpiration and thus evaporative cooling and is illuminated by a PAR lamp, so will subsequently be maintained at a slightly elevated temperature relative to the abaxial surface. This is expected to affect the *w*_sat_ estimated from bulk leaf temperature. Finally, were *w*_i_/*w*_sat_ =1, then with any symmetric uncertainty, half of the measurements would be expected to exceed unity.

Now, returning to the third and last phase of our argument, we consider whether assuming isotopic homogeneity can reproduce results on unsaturation and, qualitatively, isotopic composition. We found from the data of Wong *et al*. (2022) in Figure 7, that the isotopic homogeneity method with the maximal humidity constraint and the Cernusak *et al*. (2018) method predicted equivalent unsaturations within uncertainties. Given that Wong *et al*. (2022) concluded that their method of measuring unsaturation produced equivalent results to the Cernusak *et al*. (2018) method, we thus have agreement between three independent methods of measuring unsaturation. This strongly supports the assumption that isotopic homogeneity can successfully reproduce unsaturation results. Comparing the isotopic composition of the evaporative site when assuming isotopic homogeneity to the values of the adaxial and abaxial sides when assuming saturation (Figure 8) we found an average difference of 0.13‰, and 0.9‰, respectively. The better agreement with the adaxial surface is not surprising, given that we expect it to be closer to saturation. These average differences are less than half the standard deviation of the isotopic composition (averaging abaxial and adaxial surfaces) between leaves when assuming saturation: 1.9‰. This suggests that assuming isotopic homogeneity induces a lesser effect on measured isotopic compositions than standard leaf variance. Subsequently, we expect that under the assumption of isotopic homogeneity, the qualitative behaviour of leaf water isotopic compositions found in other studies would be reproduced. Thus, the assumption of isotopic homogeneity can reproduce results on unsaturation and isotopic composition.

On the basis of the three points of our argument, we conclude that assuming isotopic homogeneity seems to be a reasonable assumption for leaf isotope gas exchange measurements that assess the two surfaces of the leaf independently.

This conclusion has two important consequences. The first is that it indicates that unsaturation occurs in leaves even under benign atmospheric demand conditions of ∼1 kPa air saturation deficit (Figure 3). This is in contrast to past work on unsaturation (e.g. Wong *et al*. (2022)), where it was necessary to consider unsaturation only in air saturation deficits increasing from some lowest measured value. The second consequence is that unsaturation is not symmetric between the two surfaces of the leaf. Generalising from *Gossypium hirsutum* to an arbitrary leaf, we conclude (on the basis of Figures 3, 4 and 6) that the substomatal cavity of the leaf surface with the greatest stomatal resistance maintains its *w*_i_/*w*_sat_ closer to unity (saturation) than the opposite side. These consequences are summarised graphically in Figure 9.

**Figure 9:**
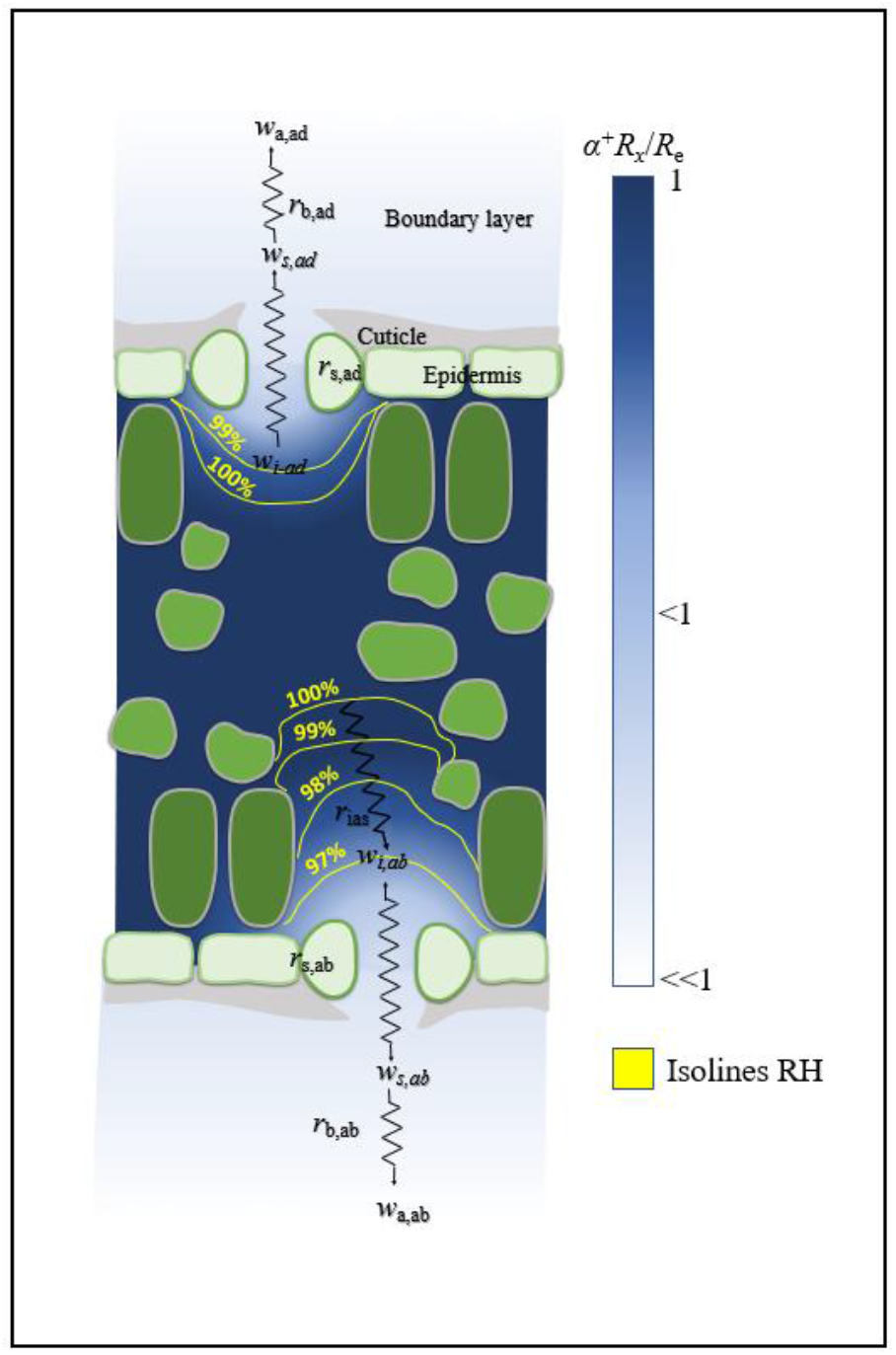
Qualitative model of a leaf at a low or mild vapour saturation deficit. Unsaturation is present, but less prominent on the surface with greater stomatal resistance (shown here as the adaxial, matching our results) and a slight isotopic inhomogeneity is generated from an internal air space resistance. For the purpose of making measurements, this isotopic inhomogeneity can be neglected, and the leaf assumed to be isotopically homogeneous.

Given that we do not consider saturation to be a reasonable assumption, we cannot comment on the spatial distribution of isotopes within the leaf; we do not necessarily expect the isotopic composition of water vapour within the substomatal cavity under the surface with greatest stomatal resistance to be depleted relative to the opposite side as found in Figure 2.

### Usefulness of the Assumed Isotopic Homogeneity Method

Before concluding, we will comment on the wider usefulness of the assumed isotopic homogeneity method. As we have argued for the reasonableness of assuming isotopic homogeneity, the application of such a method is valid and, if additional constraints are thoughtfully applied, potentially valuable to future experiments. In particular, we note that the assumed isotopic homogeneity method can be used to estimate or bound the unsaturation within the leaf, and determine its spatial extent. The determination of variability of relative humidity in the adaxial and abaxial substomatal cavities (*w*_i_/*w*_sat_) is particularly notable as it is, to our knowledge, a unique feature of our method.

With regards to which constraints to use, we consider that the maximal humidity constraint can be conceived as fixing the *w*_i_/*w*_sat_ at the surface with the greatest stomatal resistance at unity. Based on Figure 6, this seems to hold under an average *w*_i_/*w*_sat_ of approximately 90% to 100%. Subsequently the maximal humidity constraint can be used to provide a reasonable estimate of the unsaturation for benign atmospheric demand conditions. In more extreme conditions, the value generated by the maximal humidity constraint will not be expected to reflect the true *w*_i_/*w*_sat_. However, being biased towards greater humidities, it can always be used to provide an upper bound for *w*_i_/*w*_sat_. Finally, in the instance where the bulk unsaturation can be determined by another method, e.g. the Cernusak *et al*. (2018) method, then the external measurement can be used as a constraint (see equations 10-15) to determine the spatial distribution of relative humidity between adaxial and abaxial substomatal cavities.

For experiments not specifically targeting unsaturation, but instead aiming to make accurate leaf isotope measurements, determining the spatial variation of *w*_i_/*w*_sat_ using our method (as described above) is still potentially important. For instance, in our newly collected data with *Gossypium hirsutum* at 1.0 kPa air saturation deficit, ignoring leaf unsaturation produced an error between 0.13 – 0.9 ‰ (Figure 9) in the computed isotopic composition of the evaporative site. Under more extreme conditions where a greater degree of unsaturation is expected, or in a species with a greater difference between the stomatal resistances on opposing surfaces, the errors would likely be greater.

## Conclusion

This paper set out to investigate two assumptions of leaf isotope gas exchange measurements. Specifically, that the air space within a leaf is saturated with water vapour, and that the vapour is of a homogeneous isotopic composition. To this end, we presented a pair of methods for testing the feasibility of these assumptions coexisting, investigating whether assuming saturation yields the calculation of an isotopic inhomogeneity and, conversely, whether assuming isotopic homogeneity yields unsaturation. The application of these methods to data from *Gossypium hirsutum* plants at conditions of 1.0 kPa air saturation deficit and 400 ppm of CO_2_ showed significant evidence to a confidence threshold of 95% that the two commonly assumed conditions cannot coexist. Subsequently, we reasoned that an inhomogeneity exists within the leaf. To sustain such an inhomogeneity, there must necessarily be a non-negligible internal air space resistance. However, notably such an internal air space resistance would be expected to induce both variable humidity and isotopic composition, so indeed we find evidence that a real leaf is both isotopically inhomogeneous and unsaturated.

Despite this more complicated physical behaviour, we argue via a comparison with the results of Wong *et al*. (2022), that assuming only isotopic homogeneity is a reasonable approximation in the absence of a predictive model for internal resistances’ effect. In the framework of this approximation and for the case of *Gossypium hirsutum* leaves, we go on to find that the adaxial substomatal cavity remains close to saturation, even when the average leaf humidity as computed by Wong *et al*. (2022), reaches approximately 90% relative humidity. Considering this more generally, we argue that this indicates that the major unsaturation of the leaf occurs on the surface with the least stomatal resistance, and observe that this seems consistent with the bulk of transpiration occurring at this surface. Finally, we note that the method we presented for determining the unsaturation when isotopic homogeneity is assumed has potential applications in future studies of unsaturation and for making more accurate measurements of leaf isotopes at mild and high air saturation deficits.

## Author Contributions

SEC, HS-W, GDF, LAC and DAM conceived the study. SEC, HS-W, and DAM designed the experiments. SEC undertook the experimental work and data analysis. SEC, GDF and DAM developed the theory and modelling. SEC wrote the manuscript with help from all authors.

## Data Availability

All generated and analysed data from this study are included in the published article and its Supporting Information. Correspondence and requests for materials should be addressed to DAM.

## Acknowledgements

We thank Dr Chin Wong for technical assistance with the gas exchange system. We acknowledge ARC support in the form of a Discovery Grant (no. DP210103186). We thank the anonymous reviewers for their useful and valuable feedback which has improved the paper.

## Supporting Information

### Note 1: Assumed Isotopic Homogeneity Method with External Measurement as Constraint

To derive equations 11-15 from the main text, constituting the assumed isotopic homogeneity method with an external measurement as a constraint, we begin with equation 9, the parametric equation for the assumed isotopic homogeneity method without any additional constraints, and equation 10, the equation detailing the constraint imposed by an external measurement. These equations are repeated below, as equations A.1 and A.2, respectively:

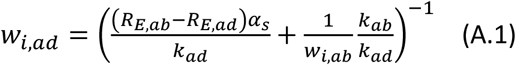

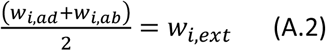

We begin our derivation by first rearranging these two equations to the forms:

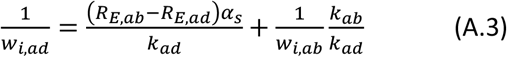

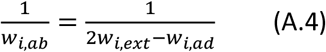

Then, substituting equation A.4 into equation A.3 yields:

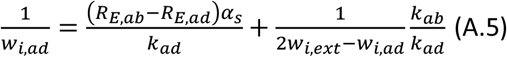

Multiplying both sides of the equation by the term *w*_*i,ad*_*k*_*ad*_(2*w*_*i,ext*_ − *w*_*i,ad*_) and collecting terms by their order in *w*_*i,ad*_ yields the following equation:

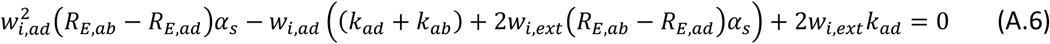

Now, we make the substitutions:

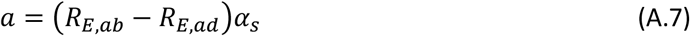

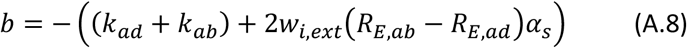

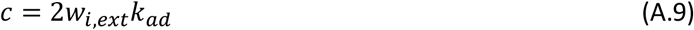

This gives us the result:

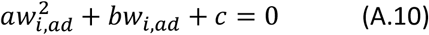

Which is immediately recognisable as a quadratic equation in *w*_*i,ad*_, so necessarily has the two solutions:

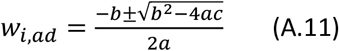

With *a, b*, and *c* as given above.

The values for *w*_*i,ab*_ corresponding to each solution can then be computed by inverting equation A.4, in particular, finding that:

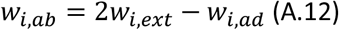

